# Consistent individual differences drive collective behaviour and group functioning of schooling fish

**DOI:** 10.1101/131094

**Authors:** Jolle W. Jolles, Neeltje J. Boogert, Vivek H. Sridhar, Iain D. Couzin, Andrea Manica

## Abstract

The ubiquity of consistent inter-individual differences in behaviour (‘animal personalities’) ^1,2^, suggests they may constitute a fundamental component of animal groups that may drive their functioning ^3,4^. Despite increasing evidence that highlights their importance ^5–16^, we still lack a unified mechanistic framework to explain and predict how inter-individual differences may affect collective behaviour. Here we investigate how differences in individual behavioural tendencies affect the group structure, movement dynamics, and foraging behaviour of animal groups using free-swimming stickleback fish (*Gasterosteus aculeatus*). Analysis of high-resolution tracking data of known individuals demonstrates that, across a range of contexts, consistent inter-individual differences in social proximity were strongly linked to speed and, together with boldness predicted spatial positioning and leadership within groups, differences in structure and movement dynamics between groups, as well as individual and group foraging performance. These effects of inter-individual behavioural variation on group-level states emerged naturally from a generic model of heterogeneous, self-organising groups. Our study combines experimental and theoretical evidence for a simple mechanism to explain variation in the emergence of structure, dynamics, and functional capabilities of groups across social and ecological scales. In addition, we show that individual performance was conditional to the group composition, providing a potential basis for social selection driving behavioural differentiation between individuals.

---

In recent years it has become apparent that across a wide range of animal taxa, individuals differ consistently from one another in their behaviour ^1,2^ (‘animal personalities’), often with large fitness consequences ^17^ and wide-ranging ecological and evolutionary implications ^18,19^. Such variation could provide a level of heterogeneity within animal groups that may drive collective behaviour. Indeed, recent studies have started to provide support for that notion, and shown that consistent behavioural differences can influence leadership ^5–8^, social network structure ^9,10^, collective dynamics ^11,12^, and group performance ^13–16^. However, rarely are consistent behavioural differences integrated within the mechanistic framework of collective behaviour research ^12,20^, which has demonstrated that relatively simple interaction rules play an important role in the emergence of collective behaviour ^21–23^. It therefore remains unclear how consistent individual differences drive the structure, movement dynamics, and functioning of animal groups.

Here, we combine high-resolution tracking of known individuals in free-swimming stickleback (*Gasterosteus aculeatus*) shoals, with agent-based models of self-organising groups, to provide a more mechanistic and predictive understanding of the behaviour, structure, and performance of groups across ecological contexts. To capture the essential dynamics within and between groups, we employ a deliberately simple, spatially explicit model, which has previously been used successfully to explain the emergence of leadership, group structure, and consensus-decision making in a range of species ^12,24–27^.

We first tested 125 fish in two classic assays to measure individual behavioural tendencies (see Figure S1). We found consistent inter-individual variation in fish’ tendency to leave a refuge and explore an open environment (repeatability R_C_ = 0.48, 95% confidence intervals: 0.33-0.60). This exploratory tendency, which may increase potential predation risk ^28^, was positively linked to fish’ food consumption, even in the safety of the holding compartment ^29^, and may reflect a willingness to accept a degree of risk in return for potentially higher foraging gains ^5^. In line with previous work ^5,7,16^ we refer to this tendency as ‘boldness’. We also found consistent individual differences in fish’ proximity to a confined shoal of conspecifics (R_C_ = 0.58, 0.46-0.68). This tendency, typically referred to as ‘sociability’ ^30,31^, was not correlated with boldness (*r*_123_ =*-* 0.05, p = 0.670), but was strongly, negatively linked with the swim speed of the fish (see Figure S1). Fish not only swam faster the further they moved from the shoal, but fish that were consistently closer to the confined shoal also had consistent higher speeds (*r*_123_ =*-* 0.79, p *<* 0.001), even in the boldness assay. This result concords with the general observation that individuals that move slower tend to form more cohesive groups ^25^, and suggests differences in social proximity may partly result from intrinsic differences in speed. We therefore prefer to refer to fish’ ‘social proximity tendency’ rather than ‘sociability’.

After quantifying the individual behavioural tendencies of the fish, we tagged all individuals for identification (see Methods) and allocated them randomly to groups of five (n = 25 groups; see Figure S1). In their natural habitat, animals may experience open, homogeneous spaces, encounter resources in spatial and temporal patches, and use habitat structures to hide from predators ^30,31^. We therefore tested the groups repeatedly in three contexts that reflect these different, ecologically-relevant scenarios, each set up in the same large, circular tanks (Figure 1A-C). Using custom-written software, we automatically identified and tracked the position of each fish in the moving groups, and computed fine-scale spatial, movement, and foraging data (Figure 1D; see Methods).

**Figure 1.**
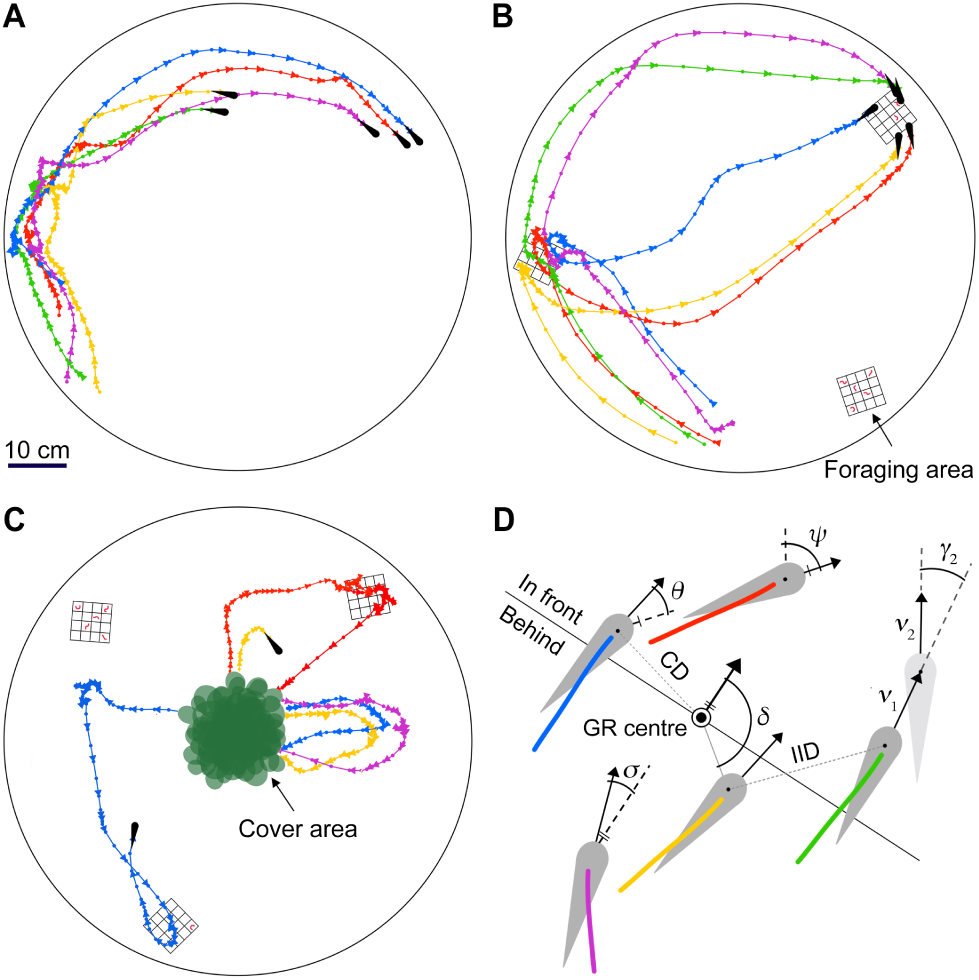
Group shoaling experiments. Schematics of (**A**) the free-schooling context, (**B**) the open foraging context with patches of food, and (**C**) the semi-covered foraging context with patches of food and plant cover. Schematics show tracking segments of one randomly selected group, with colours corresponding to the individual fish. Triangles point in the direction of motion. (**D**) Graphic illustrating key spatial and movement characteristics with arrows depicting movement vectors. For individual assays see Figure S1.

On average, sticklebacks moved in highly cohesive, ordered shoals and maintained clear zones of attraction and repulsion, mediated by relative changes in their speed and heading (Figure S2), in high accordance with other fish species ^32,33^. However, large and consistent differences existed between the structure and movement dynamics of the groups. To investigate how this variability was linked to variation in the individual behavioural tendencies within them, we employed a linear mixed modelling approach (see Methods).

We first exposed the groups to the conventional collective scenario ^23^, free movement within an open, homogeneous environment (Figure 1A). Individuals with a lower social proximity tendency, which moved faster in the individual assays, had larger nearest-neighbour distances (*χ*^2^ = 26.79, p *<* 0.001; Figure 2A) and moved more quickly (*χ*^2^ = 8.70, p = 0.009) in the dynamically moving shoals. At the same time, individual group members also strongly conformed in their speed (c.f. ^34^), a requirement to maintain cohesion, and adapted their speed depending on the mean social proximity tendency of their group (*χ*^2^ = 7.68, p = 0.012).

**Figure 2.**
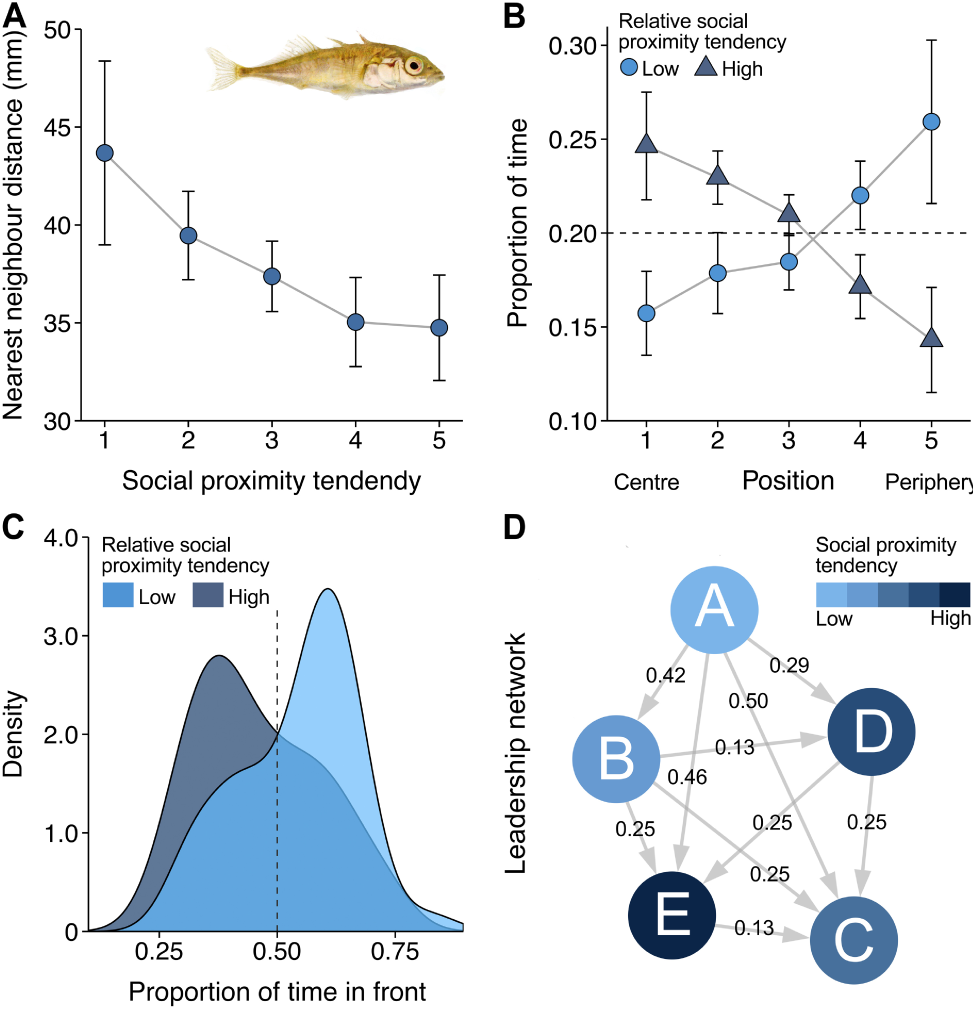
Effect of social proximity tendency on spatial positioning and leadership. (**A**) Fish nearest neighbour distance in groups as a function of their social proximity tendency, shown in 5 equally sized categories (mean *±*2 SEM; n = 120 fish). (**B**) Proportion of time fish occupied the most central to the most peripheral position in the group, calculated for each frame and averaged per individual across all frames (mean *±* 2 SEM). (**C**) Density plot of the proportion of time individuals spent in front of the group centre for the full 30 min trial. (**D**) Visualisation of a leadership network in terms of propagation of speeding changes of one randomly selected group. Numbers indicate the average temporal delay in seconds and arrows point in the direction of propagation, see Figure S3. For plots (**B**) and (**C**), individuals were evenly distributed into three categories, with the intermediate category not shown for clarity. Data were analysed as a continuous variable. See Figure S4 for model simulations.

As a result of relative differences within groups, fish with a lower social proximity tendency than their group mates, which thus tended to move more quickly, occupied positions towards the periphery (*χ*^2^ = 29.98, p *<* 0.001; Figure 2B) and front of their group (Figure 2C), an effect that strengthened over time (5 min: ΔAIC = 38.59 vs. 30 min: *χ*^2^ = 9.14, p = 0.008). This is line with recent work on pigeons, which showed that faster birds tend to lead ^8^. By assessing the propagation of movement changes in the groups ^35^, we found that such fish with a low social proximity tendency were also much more influential in deciding group motion (Figure S4), and, as a result, relative directional leader-follower networks emerged (Figure 2D). In contrast, consistent differences in boldness had no effect on the spatial positioning (*χ*^2^ = 0.64, p = 0.510) or leadership (proportion in front: *χ*^2^ = 0.06, p = 0.804) within the groups.

From the interaction of individuals with different behavioural tendencies, large differences in structure and movement dynamics emerged between the groups. Shoals with a low mean social proximity tendency predominantly schooled when together as a group (*r*_s_ = *-*0.52, p = 0.015), and moved relatively quickly, with high alignment and spacing between individuals (Figure 3). In contrast, shoals composed of individuals with a high social proximity tendency predominantly swarmed, and moved slowly, with little alignment and high cohesion between individuals (Figure 3). Measuring the strength of social interactions in the groups, we found that the strongest social forces were exhibited in the fastest moving groups (Figure S2D; c.f. ^32^). This suggest that social forces are actually weaker amongst individuals with a high social proximity tendency, which are traditionally labelled as more sociable. Boldness had no effect on these group movement dynamics (schooling: *r*_*s*_ = 0.23, p = 0.340).

**Figure 3.**
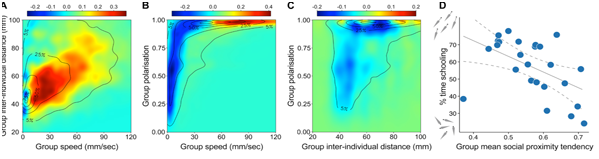
Group structure and movement dynamics in relation to group mean social proximity tendency. (**A-C**) Heat maps showing the distribution and link between the three key components of collective motion for groups with a low mean social proximity tendency (n = 13) relative to groups with a high mean social proximity tendency (n = 12). Plots are based on frame-by-frame data at time steps of 1/24th s, with groups evenly allocated to two categories based on their mean social proximity tendency. Positive values (in red) and negative values (in blue) thus indicate groups are respectively more and less likely to occupy that parameter space. Contours represent iso-levels in percentage of the highest bin for all groups combined, see Figure S2. (**D**) Proportion of time groups were schooling, characterised based on the raw distributions of group speed, cohesion, and polarisaton (see Methods). Solid grey line and dashed grey lines indicate a linear fit to the data with 95% confidence intervals.

We adapted a classic agent-based model of animal groups by incorporating variability in individual speed and goal-orientedness ^24,27^. Simulations of the self-organised, heterogeneous groups naturally result in the observed relationships, both of fish’ social proximity tendency with the spatial positioning of individuals and the structure and movement dynamics of groups, as well as the lack of such effects for fish’ boldness tendency (Figure S4).

Building on previous work ^8,25,32^, our study combines empirical data from individual and group assays with model simulations to provide evidence that individual differences in speed are a causal mechanism that drives group states, including the structure, leadership, cohesion and movement dynamics of groups. Due to relatively higher speeds, individual group members passively arrive at positions near the edge and front of groups, which in turn increases their propensity to lead. At the same time, higher individual speeds increase the speed of the group, which thereby passively results in higher order (alignment) and spacing between individuals. Differences in speed can be intrinsic as well as an emergent property, both of other intrinsic (e.g. size) and labile (e.g. nutritional state) characteristics as well as external factors (e.g. predation risk). This provides a simple candidate mechanism by which collective behaviour can emerge passively from individual differences without the need for global knowledge. Our results suggest that consistency in social proximity of the fish could, at least partly, be explained by intrinsic speed, warranting further work to investigate the extent that social proximity is actually driven by an intrinsic social tendency of individuals.

To investigate further the functional consequences that may arise from the inter-individual differences within groups, we exposed the groups to an open and semi-covered environment with patches of food (Figure 1B,C; see Methods), and analysed group foraging dynamics and performance. We found that fish that exhibited a low social proximity tendency were most likely to first discover the foraging areas in an open foraging context (Figure 4A), in line with their higher swim speed and tendency to be in the front (Figure 2C). When the environment also contained a refuge for cover, individuals spent considerable time hiding and groups often split, and now boldness best predicted what fish discovered the foraging areas (traits *×*context: *χ*^2^ = 5.77, p = 0.029). Overall, bolder fish were much faster to feed (survival model SM: *z* = 3.63, p = 0.001; Figure 4B), and more likely to initiate foraging trips in the semi-covered foraging context, thereby leading their group mates out of cover (*χ*^2^ = 8.15, p = 0.011), but also spending more time out of cover alone (*χ*^2^ = 10.28, p = 0.005; Figure 4C), a quantity that may potentially lead to higher risk of predation ^28,30^.

**Figure 4.**
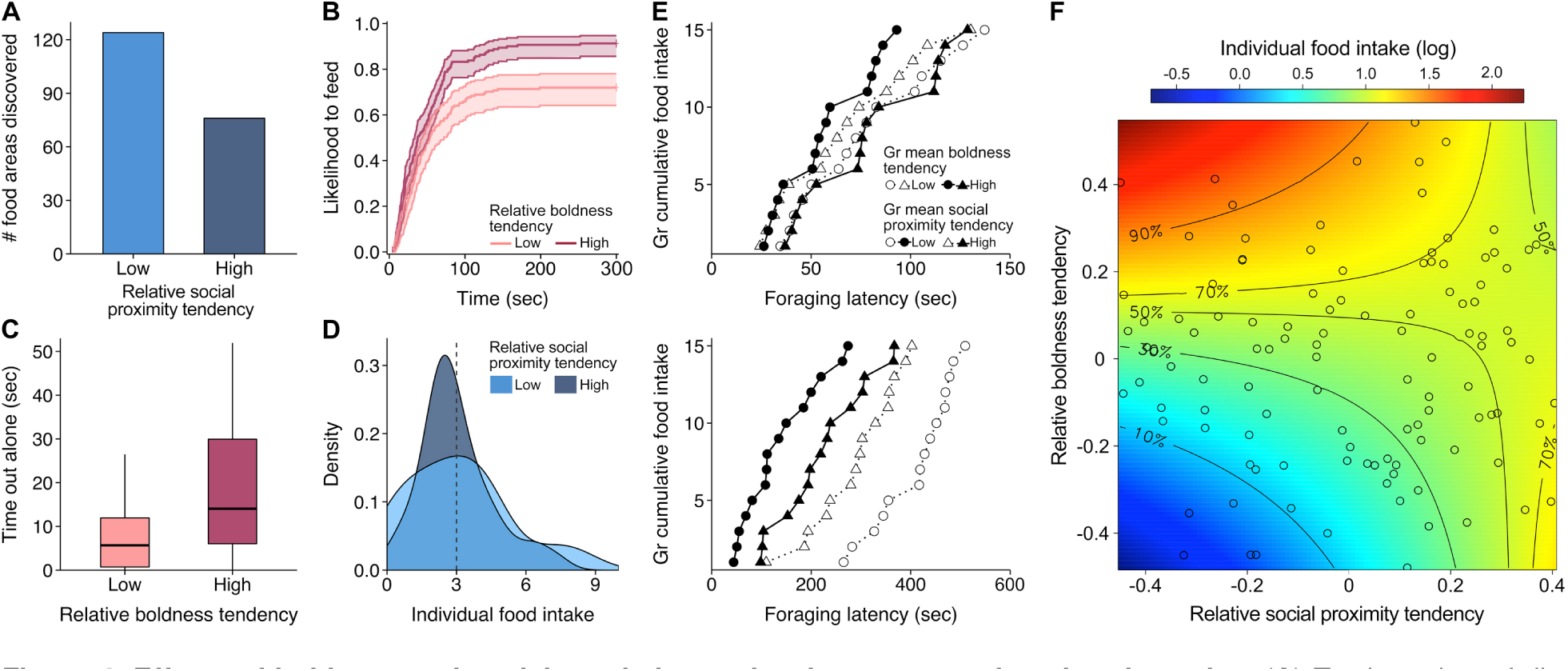
Effects of boldness and social proximity tendencies on group foraging dynamics. (**A**) Total number of discoveries during the open foraging context trials (out of 295 discoveries). (**B**) Inverted survival plot with confidence intervals of fish likelihood to feed in the open and semi-covered foraging context. (**C**) Box plots depicting total time spent out of plant cover when food was still available in the semi-covered foraging context. (**D**) Density plot of the average number of food items eaten per trial across both foraging contexts. For plots (**A-D**), individual tendencies were evenly distributed into a low, medium, and high category (n = 42, 42, 41 fish respectively), with the intermediate category not shown for clarity. (**E**) Group foraging speed in the open (upper panel) and semi-covered foraging cover context (lower panel) in terms of the latency to consume each food item (15 provided per trial). Plot shows latencies averaged across trials for each group, and groups split in four categories based on their mean boldness and social proximity tendencies (low-low; low-high; high-low; high-high: n = 5, 8, 8, 4). (**F**) Surface plot of the average number of food items eaten (log-transformed) in the open foraging context (points indicate individual fish), based on a glmm fit to the data, cropped to 90% to show the effect without fish with the extremest tendencies (n = 112 fish). For model simulation results, see Figure S4.

It was these combined effects of fish’ social proximity and boldness tendencies that ultimately explained group performance. Overall, groups composed of bold fish with low social proximity tendencies found and depleted the food patches most quickly (SM: *z* = *-*2.35, p = 0.033), with the relative effect of boldness intensified due to the availability of cover (*z* = 2.64, p = 0.016; Figure 4E). Both boldness and social proximity tendencies also predicted the foraging performance of the individual fish, relative to the mean tendency of their group (absolute differences: AIC = +13.94): fish had higher food intake the higher their relative boldness and social proximity tendency (Figure 4F), with the food intake of bolder fish enhanced by the availability of cover (traits *×* context: *χ*^2^ = 10.32, p = 0.005). Fish with lower social proximity tendencies experienced greater variance in food intake (*F*_41,39_ = 2.06, p = 0.041; Figure 4D and 4F), in line with the prediction that leadership positions come with higher variance in fitness ^36^.

Again, the general relationships of the fish’ boldness and social proximity tendency with the foraging performance of both groups and individuals, emerged naturally in simulations of our agent based model: groups with high mean speed and goal-orientedness depleted food patches the quickest and individuals with high speed and goal-orientedness had the highest food intake (Figure S4). This shows that while boldness is intrinsically linked to the foraging motivation of individuals ^5,16,29^ and thereby drives group foraging performance directly, differences in social proximity tendency have a passive effect on foraging performance by the emergent effects of speed.

In summary, we present results from individual and group experiments in combination with model simulations that provide evidence that consistent individual differences drive the spatial positioning and leadership within groups, differences in structure and movement dynamics between groups, and in turn group and individual foraging performance. This role of inter-individual differences provides a simple, parsimonious mechanism by which collective behaviour and group functioning can emerge without individuals having global knowledge of their group, providing fundamental insights that may help explain and ultimately predict the emergence of complex social and ecological patterns. We also show that individual spatial positioning and foraging performance was conditional on the composition of the group fish were in. Over time, this could result in behavioural feedback loops that may lead to behavioural differentiation between individuals via social selection, which may help explain the evolutionary maintenance of personality types ^36,37^. Our study calls for a new generation of theoretical and empirical work that further integrates individual differences with collective behaviour to better understand the multi-scale consequences of consistent individual behavioural variation, from within-group positioning to group formation and population dynamics ^37,38^, as well as its potential causal links via group-dependent individual performance.

**Supplemental Information** includes four figures and three videos and can be found with this article online.

## Acknowledgements

We thank Lucy Aplin, Damien Farine, Matt Grobis, Rufus Johnstone, Jelle Jolles, Kate Laskowski, Máté Nagy, Mike Webster, and Peter Winsberger for discussions or comments on the manuscript, and Ben Walbanke-Taylor for logistical support. We acknowledge financial support from the Biotechnology and Biological Sciences Research Council (Graduate Research Fellowship to J.W.J), the Association for the Study of Animal Behaviour (Research Grants to J.W.J and N.J.B), the Royal Society (Dorothy Hodgkin Fellowship to N.J.B), the National Science Foundation (PHY-0848755, IOS-1355061, EAGER-IOS-1251585 to I.D.C), the Office of Naval Research (N00014-09-1-1074, N00014-14-1-0635 to I.D.C), the Army Research Office (W911NG-11-1-0385, W911NF-14-1-0431 to I.D.C), the Human Frontier Science Program (RGP0065/2012 to I.D.C), the Ministerium für Wissenschaft, Forschung und Kunst 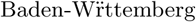 (SI-BW to I.D.C), and the Max Planck Institute for Ornithology.

## Author contributions

The study was conceived by J.W.J. and A.M with important input from N.J.B. J.W.J. and N.J.B performed the experiments, I.C. and V.S. performed the model simulations and had extensive additional input. J.W.J analysed the data. J.W.J drafted the manuscript with substantial contributions from all other authors.

## Methods

### Experimental system

We collected three-spined sticklebacks (*Gasterosteus aculeatus*) during the summer of 2014 from a stream near Cambridge, England, and housed them in our lab under controlled temperature (14 *±*1°C) and light (12h:12 h light:dark) conditions. Fish were kept in large glass tanks (120 cm length *×*60 cm width *×* 60 cm height) with artificial plants and shelters, which were maintained by both under-gravel and external filtration. Fish were fed defrosted bloodworms (*Chironomid* larvae) *ad libitum* once daily. After an acclimatisation period of six months, when fish were about nine months old, we randomly selected

125 individuals, controlling for size (body length ‘BL’: 40.6*±* 0.4 mm), and moved them to individual compartments (18.5 cm*×* 9.5 cm), each lined with gravel and containing an artificial plant. Compartments were divided from neighbouring compartments by perforated transparent partitions. We pseudo-randomly (controlling for holding tank to minimise potential familiarity effects) allocated individuals to one of 25 groups of five after the completion of the individual behavioural assays (described below). Since it is impossible to non-invasively sex sticklebacks outside the breeding season, all groups were assumed to be of mixed sex, with group sex ratio unlikely to have a big impact on our results under these controlled laboratory conditions ^39^ as both sexes are non-territorial and actively shoal together. During the whole experimental period, fish were fed three bloodworms at the end of each day. Animal care and experimental procedures were approved by the Animal Users Management Committee of the University of Cambridge as a non-regulated procedures-regime.

### Experimental overview

To control for potential social modulation and acclimatization effects ^40,41^, experiments started three days after individual housing. We started with the individual behavioural assays and subjected fish to the boldness assay on experimental days 4 and 8 and the sociability assay on days 6 and 10. We then allocated individuals to groups of five, which is a common group size for stream-inhabiting sticklebacks and conforms with previous work, which has predominantly looked at group sizes between 2-30 individuals ^7,13,32,33^. Group size and composition were kept constant throughout the experimental period. To enable individual identification in the groups, after two rest days (day 13) we tagged fish on their middle dorsal spine with a uniquely coloured disc-shaped tag (6 mm diameter) made from coloured electrical tape. This non-invasive tagging method only took between 15-30 sec per fish and has been shown to have no major effects on either the activity or shoaling behaviour of three-spined sticklebacks ^42^. After another rest day, we started with the shoaling experiments using two replicates of a large circular tank. The experimenters were blind to the identity of the fish and the composition of the groups. On day 15 we tested groups in the open tanks without food or cover, on days 16 and 17, twice per day, with patches of food but without plant cover, and on days 18 and 19 with food patches and a plant cover.

### Individual behavioural assays

Individual fish (n = 125) were tested using two standard personality assays, conventionally used to quantify boldness and sociability ^7,16,43^. The ‘boldness’assay consisted of a white Perspex tank (55 cm*×* 15 cm*×* 20 cm) containing a deep area (15 cm*×* 10 cm; 13 cm depth) with an artificial plant as refuge, and an open sandy area with a slope leading to shallow water (3 cm) at the other side (Figure S1A). We quantified the proportion of time fish spent out of the refuge as well as their average speed when at least one body length away from the refuge. The sociability assay consisted of a tank (50 cm *×* 30 cm, 8 cm depth) that was lengthwise divided by two transparent partitions to create one larger middle compartment (30 cm width), used for the focal fish, and two smaller side compartments (10 cm width), one of which contained five conspecifics (Figure S1C). At the start of each test day the fish forming the conspecifics shoal were randomly selected from the stock tanks and allowed to acclimatise to the compartment for 45 min. The position of the compartment housing the five fish was then randomly selected every four trials after which the shoal was allowed to acclimatise for 10 more minutes before the start of the next trial. In this assay we quantified the average distance from the shoal compartment as well as their speed (see Figure S1). Trials lasted 30 and 15 minutes for the boldness and sociability assay respectively. For both assays, fish were taken from their individual compartment at the start of a trial and returned there immediately after completing the trial using a dip net. We used a custom replicated set-up of eight boxes that enabled us to test multiple fish simultaneously under identical conditions, while minimising outside disturbances. Sessions were automatically recorded at 12 fps in high-definition using Raspberry Pi computers (Raspberry Pi Foundation, England) positioned in the top of each box.

### Group shoaling experiments

To investigate the collective behaviour of the fish, groups were repeatedly subjected to a white, circular Perspex tank (80 cm diameter, 20 cm height; 7 cm water depth), positioned inside a large white light tent (200 cm *×* 100 cm *×* 160 cm) illuminated from all sides. For the fish, the tank is a potentially dangerous environment due to being bright, open, and homogeneous, and results in fish to strongly school together (see Figure S2). For the trials in the foraging contexts we placed three food patches at random locations in roughly equilateral triangular formation in the tank, between 5 cm from the wall and 15 cm from the tank centre. Food patches consisted of white Perspex grids (5 cm *×*5 cm *×*1 cm) containing five bloodworms each, randomly distributed among the grids’ 16 cells. The patches were constructed such that fish would notice the prey items from a distance of approximately 10-15 cm. For the trials in the semi-covered foraging context, artificial plants were positioned in the centre of the tank, creating a covered area with a diameter of 15 cm.

Each group received a total of seven test trials: one in the classic context (30 min), four in the open foraging context (5 min), and two in the foraging plus cover context (10 min). The group order of testing was randomised but a fixed context order was used as not to confound the behaviour of the fish in the earlier contexts with experience of the foraging patches and cover being available. Data analysis of the free-schooling context trials focused on the first 5 min only (c.f. ^33^) but trials lasted 30 minutes to enable the analysis of certain temporal effects (see below). Before each trial, fish were taken from their individual compartment using a dip net and allowed to acclimatise for 30 sec in black plastic cups, after which all five fish of a group were simultaneously placed in a transparent Perspex cylinder (10 cm diameter) in the centre of the tank. After another 30 sec acclimatisation, the fish were released by remotely raising the cylinder. At the end of each trial, fish were placed back in their compartments, potential waste and remaining food items removed, and tank water circulated to mix any chemical cues. Trials were recorded from above at 24 fps at a resolution of 1400 *×*1400 using Raspberry Pi computers. As groups received two foraging trials per day, with 5 bloodworms provided in each foraging patch, hypothetically a fish could reach a maximum daily food intake of 30 food items. This was by far never observed. Furthermore, sticklebacks under similar conditions are capable of consuming up to 60 bloodworms within a three-hour timespan ^29^. Satiation is therefore unlikely to have had a strong effect on the observed foraging performance.

### Automated tracking and data collection

We acquired highly detailed individual-based movement data for both individual and group assays with custom tracking software written in Python version 2.7.12 (by J.W.J.) using the OpenCV library. For the individual trials, a background image, created by averaging the first 200 frames, was subtracted from each frame and fish identified via automatic tresholding using constant threshold values. For the group trials we automatically identified fish based on their differently coloured tags, which enabled us to acquire highly accurate tracking data linked to each individual, despite occasional occlusions. Positional coordinates were converted from pixels to mm and subsequently smoothed using a Savitzky & Sgolay smoothing filter with a window of 15 frames. After tracking, all trajectory data were visually checked for any inconsistencies or errors and, if needed, manually corrected. In addition, we performed manual video observations for the trials in the foraging contexts and recorded the time each food item was eaten (0.1 sec precision) as well as the identity of the foraging fish.

### Computation of behavioural data

#### Individual characteristics

We determined each fish’s velocity, speed, direction, acceleration, and turning speed directly from the discrete tracking data using the following series of calculations. With the vector **r**_*i*_(*t*) = (*x*_*t*_(*t*)*, y*_*i*_(*t*)) denoting the position of fish *i* at time *t*, we approximated its velocity **v**_*i*_(*t*) = (*u*_*t*_(*t*)*, w*_*i*_(*t*)) using the forward finite difference

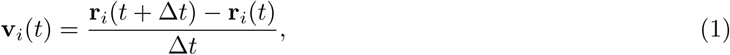

where Δ*t* = 1*/*24 s is the time interval between subsequent position measurements. The speed *v*_*i*_(*t*) is then given by the norm of the velocity vector, such that

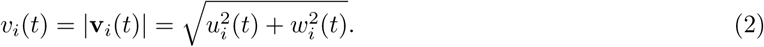

Next, we quantified the direction of motion using the angle *ψ*_*i*_(*t*) between the velocity vector and the positive *y*-axis, which is given by

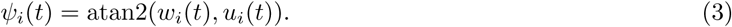

Furthermore, we quantified the acceleration as a finite difference of the velocity

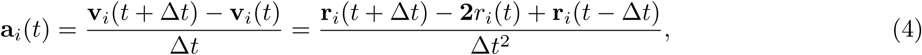

and the turning speed, or angular velocity, as a finite difference of the angle,

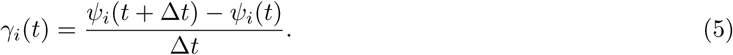

As fish were placed at the origin of the Cartesian coordinate system pointing north, care was taken to compute the correct angular difference with regard to the periodicity of *ψ*_*i*_(*t*)

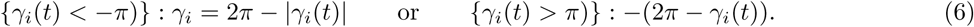

#### Within group positioning

We determined the positioning and ordering of the fish in a group relative to one another and to the direction of motion of the group centre using the following calculations and linear transformations. To calculate fish nearest neighbour distance (NND), we computed a matrix of distances between all individuals and then determined the minimum value for each fish and calculated the distance as

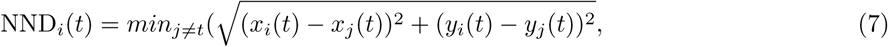

where (*x*_*j*_(*t*)*, y*_*j*_(*t*)) denotes the coordinates of the *j*th neighbour.

Next, for each time step we identified the mean coordinates of all fish in a group 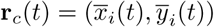, that is, the group centre, and then estimated the velocity *v*_*c*_(*t*) and direction *ψ*_*c*_(*t*) of the group centre at time *t* using the calculations as for the individual fish (described above). Then for each frame we calculated the distance of each fish to the group centre as

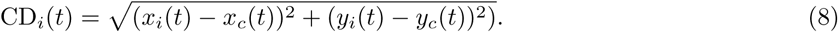

To calculate relative positions of individuals to the group, we shifted the coordinates of each fish so that the origin of the coordinate system was at the group centroid, and determined the angle between the positive *y*-axis through the group centroid and an individual’s position

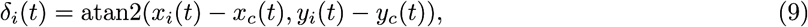

which we subsequently used to calculate an individual’s relative direction to that of the group centre

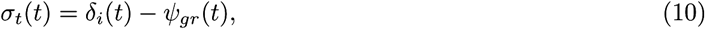

which we then adapted to fit to the Cartesian coordinate system pointing north

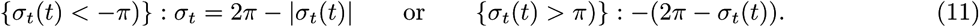

Based on the relative direction and distance to the group centre, we then shifted the coordinates of each fish so that the origin of the coordinate system lay at the group centre, and its direction pointed north, aligned with the positive *y*-axis:

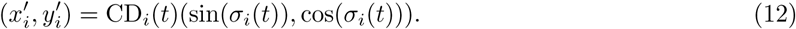

The transformed coordinates of the fish meant that fish with greater *y*-coordinates were at the front for a given time step. We then counted the proportion of frames that each fish was located in front of the group centre. To further examine inter-individual positioning in the group, we calculated fish’ relative direction to that of its four group mates *θ*_*ij*_ from the respective angles of the fish with the *y*-axis (*ψ*_*j*_) following the calculations as used for the relative positioning to the group centre.

#### Group characteristics

To examine the properties of the differently composed groups, we calculated the speed of the group centre, group cohesion, and polarisation using the following calculations. For each time step *t*, the speed of the group *v*_*c*_(*t*) is given by the norm of the velocity vector, such that

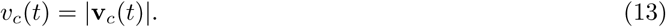

We then calculated the average inter-individual distance IID_*c*_(*t*), as a measure of group cohesion, based on the individual distances IID_*ij*_ between all fish in a group

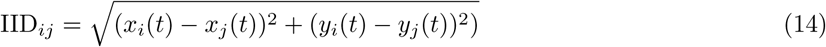

using

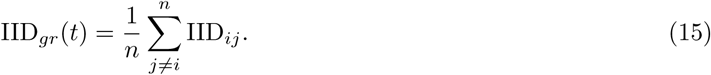

And finally we calculated the polarisation of the group

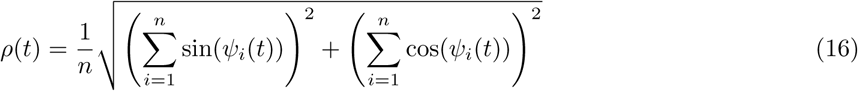

which is a measure of the alignment of the fish in the group relative to each other, and ranges from 0 (complete non-alignment) to 1 (complete alignment).

Schooling is defined as a cohesive group that moves with considerable speed and alignment, while a group is said to swarm when it is cohesive but has no or little speed and/or alignment between its members ^21^. To investigate the schooling tendency of the groups, we computed the distributions of the three fundamental components of schooling on the full dataset: group cohesion, speed, and polarisation (see Figure 3 and Figure S2). Furthermore, based on the detailed distributions of all groups and parameters from previous work ^25,32^, we also categorised groups to school, based on the following criteria: average inter-individual distance IID_*gr*_ *≤* 160, speed of group centre vector *v*_*gr*_ *≥*0.5 BL/s, polarisation *ρ* ≥ 0.6, no outliers or group split. Outliers and group splits were computationally identified based on a non-linear distribution of ordered distances between all group members in terms of the IID and NND, with parameters identified based on the raw data distributions.

#### Propagation of motion

To investigate leadership in terms of the propagation of movement changes in the group, we examined temporal correlations in speeding and turning changes for all dyads within a group ^32,35^. We compared the speed and direction of the two fish in a dyad up to 72 frames (3 sec) earlier and later, in time steps of 1/24th s, and quantified the mean time point of the maximum correlation coefficient (see Figure S3). A leading event was said to have occurred when a fish’ change in speed or direction was ‘copied’ by another fish delayed in time. Subsequently, we constructed leadership networks based on the time delays between all group members following Nagy et al ^35^. Analysis was restricted to frames in which fish were less than four body lengths apart and moved faster than 1 BL/s.

#### Foraging and hiding behaviour

For the trials in the two foraging contexts we used the positional data to compute the order that individuals arrived in the vicinity (≤30 mm) of and above the foraging patches. We defined the first fish to ‘discover’ a foraging patch as the one that first arrived in its vicinity during a trial. For the trials in the semi-covered foraging context we also calculated the proportion of time individuals spent out of cover (with at least half their body), the proportion of time individuals spent out of cover alone, and their average order number for leaving cover. In turn, these measures were used to calculate the average number of fish out of cover and the proportion of time all fish were out of cover.

### Data analysis

Data were analysed in R 3.2.0. We used a generalised linear mixed modelling approach ^44^ to investigate the effects of inter-individual behavioural differences on behavioural repeatability as well as individual and group shoaling and foraging behaviour. To assess individual behavioural consistency, we calculated Consistency Repeatability ^45^ using linear mixed models that included day as a fixed effect and fish ID as a random factor. We calculated 95% confidence intervals of repeatability by running 10,000 permutations of each test. Significant effects are those with a confidence interval that does not overlap 0. Boldness and social proximity scores were scaled between 0 and 1, with social proximity values square-root transformed and inverted before scaling. Neither fish’ boldness tendency nor social proximity tendency was significantly correlated with body size (p = 0.804 and p = 0.765). The randomized group compositions (n = 25 groups) were normally distributed in terms of the average personality types.

For the behaviours in the shoaling experiments, response variables were calculated based on the the distribution of the data on a frame-by-frame basis, with mean values calculated for approximately normal (transformed) distributions and median values when data was skewed. For the individual-level models we included individual boldness and social proximity scores and the interaction between them as fixed effects. Group identity was fitted as a random factor to account for the non-independence of individuals within a group, and individual identity nested in group identity was additionally included for the trials in the two foraging contexts to account for the repeated measures-nature of the data. For the group-level models we fitted the mean boldness and mean social proximity tendency of the group and the interaction between them. We only included measures of group variability in behavioural tendencies in the case of clear a priori hypotheses as not to over-parameterise our models. Food intake and the likelihood to discover the foraging patches were fitted to a Poisson error distribution with log link function, appropriate for count data. To investigate how the effect of inter-individual differences on the proportion of time fish spent in the front of the group changed over time, we compared models based on the first 5 min and full 30 min of the trial. To investigate the propagation of speeding and turning changes in the groups, we ran an ordinal logistic regression with individual boldness and social proximity tendency ranks in the group as fixed factors, and the random data structure as described above. We analysed the foraging behaviour of individual fish and the groups over time with Cox proportional hazards (survival) regression models. Survival analyses avoid censoring the data, thereby allowing for the assumption that fish or groups assigned to maximum time may have foraged or finished all the food respectively had the trials run longer. For these analyses, the data were clustered around fish identity and group identity to account for dependence in the data and for trial to account for changes over time.

Minimal adequate models were obtained by backward stepwise elimination following Crawley ^44^, i.e. sequentially dropping the least significant terms from the full model, until all terms in the model were significant (all interaction terms were non-significant unless documented). Statistics for non-significant terms were obtained by adding the term to the minimal model. We also report AIC when comparing models when based on different subsets of the data. Residuals were visually inspected to ensure homogeneity of variance, normality of error and linearity where appropriate. Differences in variance were analysed using a Levene’s test, making sure there was no difference in variance in the personality composition of those groups. Data were log- or square-root transformed if assumptions were violated, or, where appropriate, a robust Spearman rank correlation test was used. We had to exclude one group onwards from the 4th open foraging context trial due to the death of one fish, and one trial in the open foraging context and one trial in the semi-covered foraging context due to experimental errors. For two cover context trials no foraging data could be collected due to a recording error. To control for multiple testing, we employed a False Discovery Rate correction for all statistical tests, with corrected p-values stated in the text. p *<* 0.05 is reported as significant and means are quoted *±* SEM throughout unless stated otherwise. Other statistical parameters are reported in the main text and figure legends.

### Individual-based modelling

#### Overview

We adapted a simple spatially-explicit self-propelled particle model detailed in Couzin et al ^24^ and combined it with goal-oriented behaviour (omega) ^26,27^, which showed it to be an important factor in individuals’ responses to known resource locations. We deliberately chose this simple model, not to obtain a quantitative comparison to the experiments, but to determine if the general results are consistent with theory, and seek a parsimonious explanation for the observed patterns.

#### Framework

Groups were composed of individuals, each characterised by a position vector *c*_*i*_(*t*), a unit direction vector 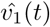 and speed *|v*_*i*_(*t*) *|*, where *i* is the identity of the individual and *t* is the current time step. The speed of each individual is drawn from a normal distribution to represent consistent inter-individual differences. Hence, each individual differs in speed and a given individual’s speed remains constant within a simulation. While having a constant speed is an oversimplification (to obtain the simplest possible model formulation that can explain the experimental results), due to the nature of response to social interactions, individuals can effectively slow down, or speed up, by virtue of modifications to the small-scale tortuosity of their motion. For example, fast individuals at the front of groups will tend to be attracted to those behind, resulting in them taking a more tortuous path, effectively slowing them in the direction of travel of the group as a whole, whereas slower individuals trailing groups will exhibit highly directed motion that increases their relative speed in the direction of travel with respect to other group members (see Video 3).

Social interactions with others were accounted for through three types of interactions: repulsion, alignment and attraction. Individuals turn away from *n*_*r*_ neighbours encountered within a small radius (*r*_*r*_) around them. This represents collision avoidance and maintenance of personal space expressed by the agents, and, as is apparent in real schools ^30,31^, takes highest priority.

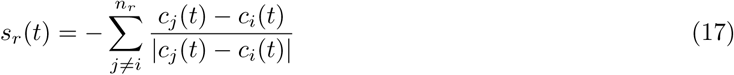

where *s*_*r*_(*t*) represents the social component of an individual’s desired direction of motion after responding to individuals within *r*_*r*_.

If no individual is present within radius *r*_*r*_, the focal individual orients itself with individuals within *r*_*a*_.

These zones are circular, with a blind area of *α*° behind the individual. In these zones, individuals interact with conspecifics only in the remaining (360 *- α*)°. All three zones are non-overlapping and their widths are defined as Δ*r*_*r*_ = *r*_*r*_, Δ*r*_*o*_ = *r*_*o*_*- r*_*r*_, and Δ*r*_*a*_ = *r*_*a*_*- r*_*o*_. Since we simulated a group of five individuals, and due to the relatively small environment in which experiments were conducted, where individuals can readily see others at the maximum possible spacing, we set the maximal range of perception, *r*_*a*_ to ∞. Each individual attempts to align its direction of motion with *n*_*o*_ neighbours in the zone of orientation, giving

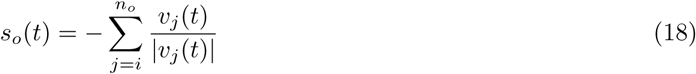

and is attracted towards positions of individuals within the zone of attraction

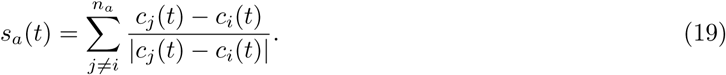

Once individuals have a social vector, they reconcile this with their goal-oriented tendency *g*_*i*_(*t*) weighted by a continuous term *ω*, which represents the strength of individual goal-orientedness. Like speed, individual *ω* is drawn from a Gaussian distribution (to represent consistent inter-individual differences) and remains constant in a given simulation

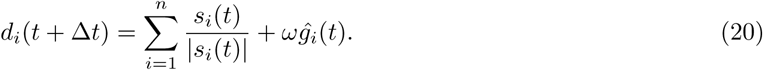

*d*_*i*_(*t* + Δ*t*) is then normalised 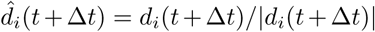, to represent the desired direction of motion of the individual. Individuals’ goal-oriented vector *g*_*i*_(*t*) points in the direction of their current motion until they enter a radius *r*_*c*_ of a rewarding cue. This can be interpreted as their inertia, or their desire to continue moving in their current direction when reward is not perceived. Once individuals are within this set radius, their goal-oriented vector *g*_*i*_(*t*) is directed towards the reward to an extent determined by their *ω*. Once individuals are on a food patch, they feed with a feeding rate *f*.

Motion of all individuals is subject to noise (error in movement and/or sensory integration) which is implemented by rotating 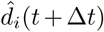 by a random angle chosen from a circularly wrapped Gaussian distribution centred at 0 and of standard deviation *e*. Once the desired direction has been determined, individuals turn towards 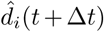 with a maximum turning rate of *ψ*Δ*t*.

In the foraging context (see below), boundary conditions were enforced by modifying the desired direction of an individual to equal a boundary vector *b*_*i*_(*t*) when they reached a narrow zone near the edge of the arena. Boundary vector *b*_*i*_(*t*) is a unit vector pointing towards the centre of the arena. This was done to allow agents to avoid walls and to prevent them from leaving the arena. In the free schooling context, individuals were initialised in a periodic boundary environment to ensure that no boundary related artefacts are observed while measuring spatial positioning of individuals.

#### Simulations

In line with the experiments, we started with simulations of groups composed of five individuals. To simulate the free-schooling context and open foraging context presented in the experiments, we initialised the groups both in an open, boundary free environment and in a circular environment that contained three food patches (10 units radius). Individuals were initialised with random positions and directions in the middle of the arena, again in line with the experimental procedure. Details about model parametrisation can be found in Table 1. Parameter values for the schooling models are standard values, previously explored in ^24,25^. To explore further how the effects may be group-size dependent, we ran additional simulations with larger groups of twenty. As speed distributions are often right-skewed and bound at zero, including our experimental data (see Figure S1), we also ran simulations of the free-schooling context (for one specific p<u>aram</u>eter condition) with a Gamma distribution of shape parameter (*k* = 0.4 and scale parameter, *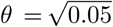*). These parameters were chosen so that the distribution had a mean within our tested range and variance identical to the one used in case of the Gaussian distribution. For the free-schooling context we ran simulations of 2,000 time steps and for the foraging context 10,000, with data being stored every 200 and 500 time steps respectively, with 400 replicates of each parameter condition explored.

**Table 1.** Summary of individual-based model parameters.

| Parameter | Symbol | Values explored |
| --- | --- | --- |
| Arena size | $A$ | 500 |
| Zone of repulsion | $r_r$ | 1 |
| Zone of orientation | $r_o$ | 6 |
| Zone of attraction | $r_a$ | ∞ |
| Field of perception | $\alpha$ | $270^\circ$ |
| Turning rate | $\psi$ | $60^\circ$ |
| Speed | $\vert v\vert $ | 0.1 – 2.0 |
| Speed error | $e_s$ | 0.1 |
| Omega | $\omega$ | 0.01 – 0.1 |
| Omega error | $e_\omega$ | 0.01 |
| Timestep increment | $\Delta_t$ | 0.1 |
| Cue detection radius | $r_c$ | 30 |
| Nr of food patches<sup>a</sup> |  | 3 |
| Nr of food items per patch<sup>a</sup> |  | 50 |
| Feeding rate<sup>a</sup> | $f$ | 0.001 |

<sup>a</sup>*Foraging context simulations only*

## Supplemental Information

**Figure S1.**
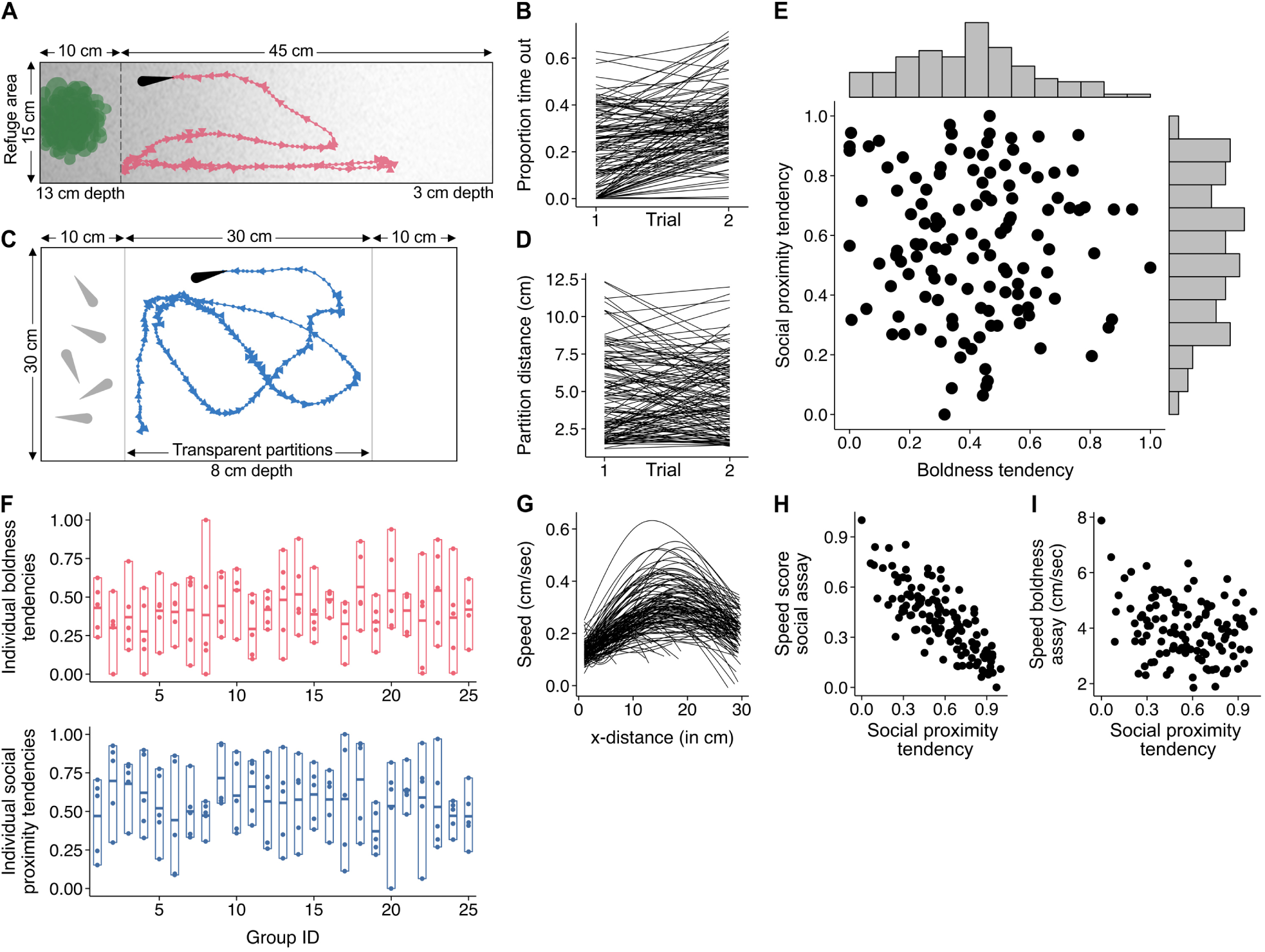
Consistent inter-individual behavioural tendencies. Related to Figure 1. (**A**) Schematic of the ‘boldness assay’, a rectangular tank with a deep refuge area that leads to an increasingly shallow open area on the other side. (**B**) Line plot showing individual repeatability in terms of the proportion of time fish spent out of the refuge (‘boldness tendency’), which was strongly, positively linked to their average distance out of cover (*r*123 = 0.668, p *<* 0.001). (**C**) Schematic of the ‘sociability assay’, a tank with a large centre compartment for the focal fish, and two side compartments, one empty and one containing five conspecifics. (**D**) Line plot showing individual repeatability in terms of the average distance from the conspecifics’ compartment (‘social proximity tendency’). (**E**) Relationship between individual boldness and social proximity tendencies (n = 125 fish) and their distributions (grey bars), with personality scores scaled between 0 and 1. These plots show that fish were highly repeatable in their tendency to explore out of cover, as well as in their propensity to stay near the confined shoal in the sociability assay, with no link between them. (**F**) Group compositions in terms of the individual group members’ boldness and social proximity tendencies (n = 25 groups of 5). (**G**) Line plot showing predicted speed curves (quadratic fit) for both testing trials of each fish in the sociability assay in terms of the distance from the compartment holding the shoal. A speed score was calculated for each fish by determining the speed where the curve was maximal, averaged across both trials, to minimise issues with restricted speed near the partition. (**H**) Relationship between fish’ social proximity tendency and their average speed score (scaled). (**I**) Relationship between social proximity tendency and average swim speed in the boldness assay (*r*_118_ = *-*0.28, p = 0.005). Plots (**G-I**) indicate a strong, negative relationship exists between social proximity and swim speed.

**Figure S2.**
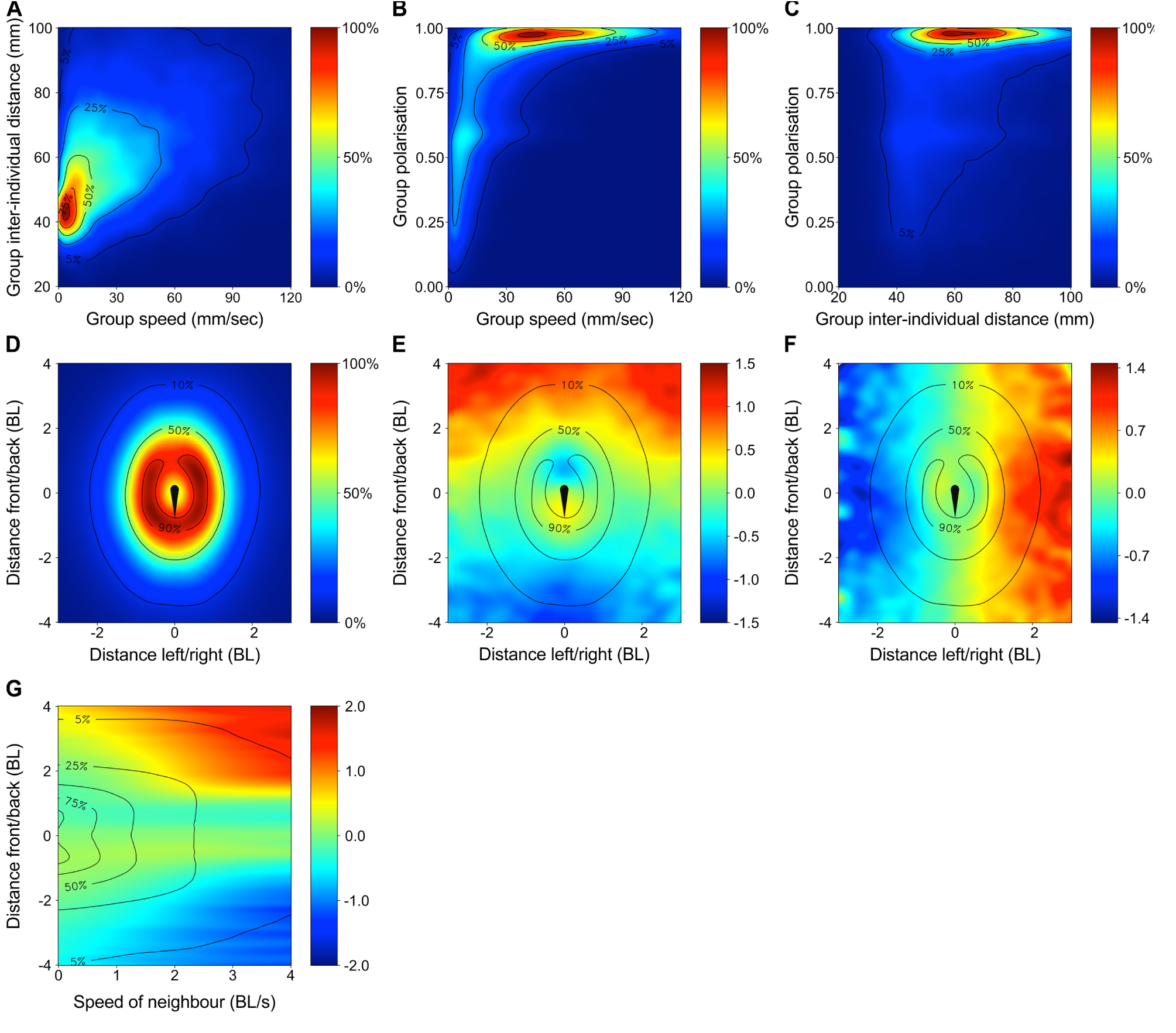
Surface plots of group and individual movement dynamics. Related to Figure 2 and 3. Plots are based on the full dataset in the open, homogeneous context with time steps of 1/24th s. (**A**) Relationship between group speed and cohesion. (**B**) Relationship between group speed and polarisation. (**C**) Relationship between group cohesion and polarisation. Data was cropped to show the most relevant area only (respectively 86.3%, 88.2%, and 97.3% of the full parameter space). Contours represent iso-levels in percentage of the highest bin for data of all groups combined. These plots indicate a strong link between group cohesion, speed, and polarisation, with faster moving groups being less cohesive and more strongly aligned. Groups moved at a steady median pace of 30.0 mm/sec, with an average group cohesion (IID) of 70.7 mm. In the direction of motion, groups had an average length of 100 mm and rarely exceeded 300 mm. Groups were strongly polarised the majority of the time (median = 0.92) and had very low levels of fragmentation, with significant outliers or group splits (for explanation, see Methods) only occurring 4.9 *±* 1.5% of the time. (**D-G**) To investigate the individual interaction rules, we selected each fish in each group and computed its position, acceleration, and turning forces relative to the position and speed of its group mates (see Methods). Distances are expressed in units average body length (‘BL’). (**D**) The probability of finding neighbouring fish at a given position relative to the position of the focal fish, which was placed at the origin pointing north. Fish density is presented in percentages relative to the densest bin for all groups combined. (**E**) and (**F**) Respectively the acceleration and turning speed of the focal fish as a function of the position of its group mates. (**G**) Focal fish’ acceleration forces as a function of the swim speed of its group mates and its front-back distance. For the turning speed, positive values indicate a right turn and negative values a left turn. Data was cropped to show the most relevant area only (**D-F**: 92.1% and **G** 93.3% of the full parameter space). These plots indicate that on average, (**D**) fish are very likely to be within one body length of another group member side-by-side, and within two body lengths front-to-back (NND = 39.0 mm), (**E**) fish speed up when a neighbouring fish is far ahead or just behind them, but slow down when a neighbouring fish is far behind or just in front, (**F**) fish turn left when a neighbouring fish is on its far left side and turn right when its neighbour is on its far right side, with weaker opposite turning tendencies when neighbouring fish are very close, and (**G**) fish acceleration forces become stronger the faster the neighbouring fish is moving.

**Figure S3.**
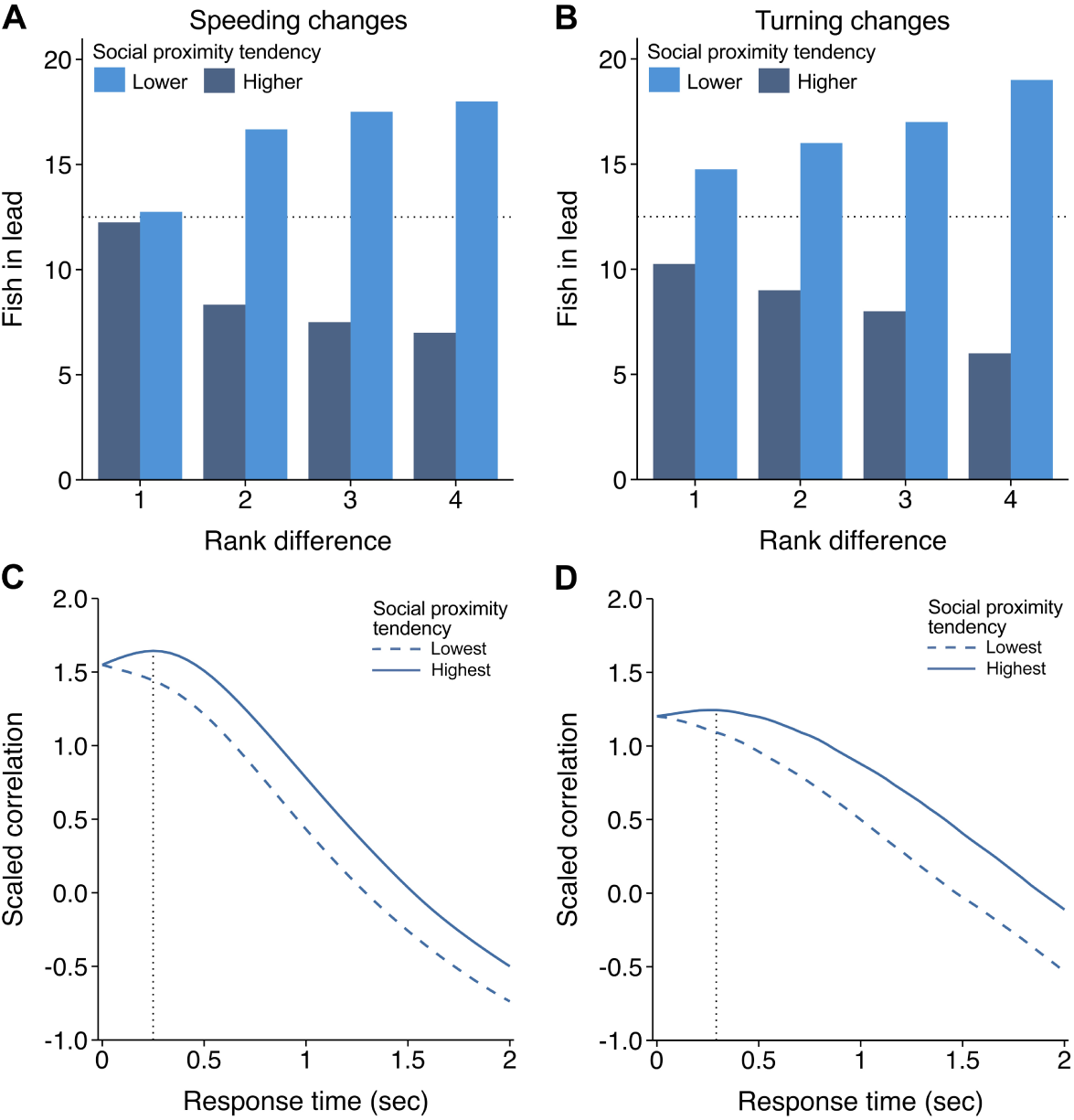
Propagation of movement changes. Related to Figure 2. To investigate the propagation of movement changes, we selected each fish in each group as focal individual and compared its swim speed and direction to that of all its group mates up to 3 seconds later, at time steps of 1/24th s. We then determined the average time difference for the highest correlation across the trial for all dyads in each group. Fish were ranked based on their social proximity tendency (rank 1-5) within each group. (**A**) and (**B**) Bar plots depicting number of dyads for which movement changes on average propagated from the fish with the higher social proximity tendency versus from the fish with the lower social proximity tendency. Bars show mean values for rank difference of 1-3 (n = 100, 75, 50 respectively) and total number for a rank difference of 4. Dotted line represents the value that both personality ranks would lead equally. (**C**) and (**D**) Median correlations in movement changes for the fish with the highest social proximity tendency relative to fish with the lowest social proximity tendency in each group and the other way around. Correlation coefficients were scaled for each group to control for between-group variability, and analysis was restricted to frames in which both fish were moving at a speed of at least 10 mm/s during the full 30 min trial in the free-schooling context. Both the (**C**) the swim speed correlation and the (**D**) turning correlation of fish with the highest social proximity tendency in a group peaked after zero, with a delay time of less than 0.5 s before decaying (indicated by the grey dotted line), whereas for fish with the lowest social proximity tendency the correlation curve does not show such a peak. This suggests that fish with a higher social proximity tendency on average speed up, slow down and turn in response to the speed and direction of fish that have a relatively lower social proximity tendency. Both speeding, *r*_123_ = 0.65, p *<* 0.001, and turning changes, *r*123 = 0.54, p *<* 0.001, were positively linked with the tendency to be in front. These plots thus show that fish with a higher relative tendency for social proximity, which moved faster in the solitary and group assays and were more in front, are more likely to lead their group mates in terms of both (**A**) the propagation of speeding changes (ordered logistic regression: *z* = *-*2.78, p = 0.012) and (**B**) the propagation of turning changes (*z* = *-*2.76, p = 0.012), and that this increases the larger the rank difference between the two fish.

**Figure S4.**
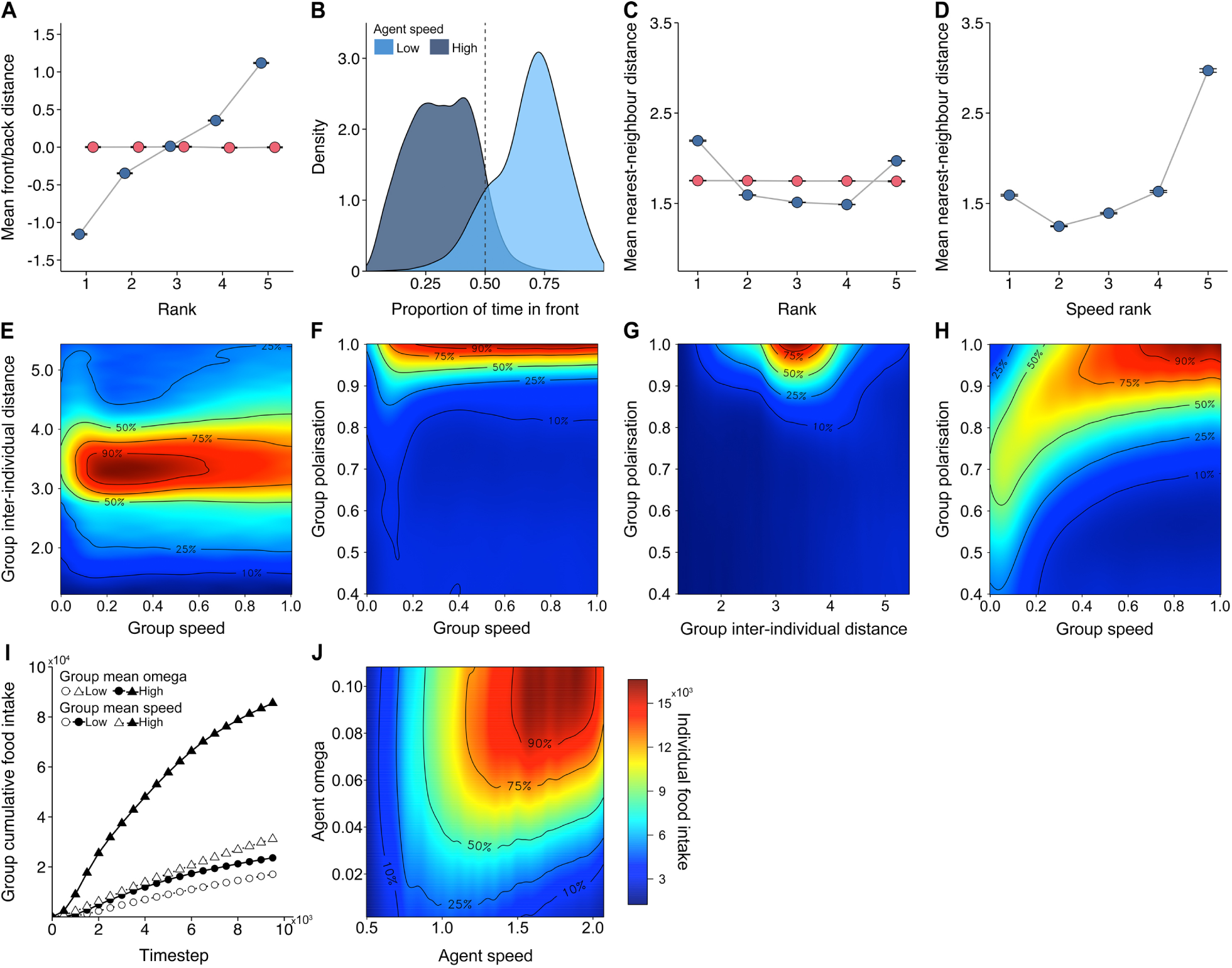
Data from the individual-based model simulations. Related to Figures 2, 3 and 4. (**A**) Mean distance in front/behind the group centroid in terms of an individual’s speed rank (blue) or omega rank (goal-directedness; red) in the group. (**B**) Density plot of the proportion of time individuals spent in front of the group centroid in terms of their set speed, categorized in three equally sized bins with the intermediate bin not shown for clarity. (**C**) Mean nearest-neighbour distance in terms of an individual’s speed rank (blue) and omega rank (red). (**D**) Mean nearest-neighbour distance in terms of an individual’s speed rank but now with speed scores drawn from a gamma distribution, see Methods and Figure S1. Front/back and nearest-neighbour distances were averaged across the simulation and expressed in units of repulsion radius. Error bars indicate 95% confidence intervals of the mean. These plots indicate that within a group, faster individuals tend to be towards the front of the group and further from their neighbours (note, also very slow individuals tend to be further away), especially when the distribution of individual speeds is right-tailed. Individual goal-directedness (omega) had no effect on these behaviours. Compare with Figure 2 and see Figure S1. (**E-H**) Surface plots depicting the relationship between (**E**) group speed and cohesion, (**F**) group speed and polarisation, (**G**) group cohesion and polarisation for groups of 5 individuals, and (**H**) group cohesion and polarisation for groups of 20 individuals. Plots are based on the full dataset but cropped to show the most relevant area only (respectively 90.3%, 90.4%, 92.2% and 85.8% of the full parameter space). Colour scale is square-root transformed and reflects z-scores in percentage relative to the highest bin, with contours representing iso-levels. Plots (**A-H**) are based on 400 replicates of 2,000 time steps taken at intervals of 200 time steps. Plots (**E-H**) indicate that faster groups, i.e. those composed of individuals with higher set speed, were sparser and more polarised than their slower counterparts. The link between speed and polarisation becomes especially clear when the group is larger, with groups of 20 needing higher speed to reach the same level of polarisation. The effect of speed on inter-individual distance, however, is weak. This is partly due to the three zone model, which allows for stable existence of neighbours in the alignment/orientation zone alone (see Methods). Compare with Figure 3 and Figure S2. (**I**) Cumulative food intake over time, showing mean values for groups evenly split into four categories based on their average set speed and goal-directedness (omega). (**J**) Surface plot showing individual food intake calculated as the number of food particles consumed by an individual in terms of its speed and goal-directedness (omega). Data was cropped to show the most relevant area (73.6% of the full dataset). (**I-J**) are based on 400 replicates of 10,000 time steps taken at intervals of 500 time steps. These plots indicate an interaction between agent speed and omega drove both group and individual foraging performance: groups depleted the food more quickly the faster and more goal-oriented they were, and within groups, individuals that were faster and more goal oriented consumed more food. This is linked to the fact that faster individuals are more in front and therefore arrived at rewards sites sooner than their group mates, while omega determined an individual’s directedness towards the food once within the cue detection radius (see Methods). Compare with Figure 4.

## Supplementary videos

**Video S1. Individual personality assays.**

Video showing the tracking of an individual fish in the ‘boldness’ assay and in the ‘sociability’ assay with the behavioural measures automatically acquired.

**Video S2. Group shoaling experiments.**

Video showing a group of fish tested in the three assays used for the group experiments: an open, homogeneous environment, an open environment with patches of food, and a semi-covered environment with patches of food.

**Video S3. Individual-based simulations of self-organising, heterogeneous groups.**

Video showing a visualisation of the individual-based simulations of self-organised groups consisting of 5 and subsequently 20 agents, with the emergence of leadership plotted dynamically over time.

## References

1. Réale, D., Reader, S. M., Sol, D., McDougall, P. T. & Dingemanse, N. J. Integrating animal temperament within ecology and evolution. Biol Rev 82, 291–318 (2007).

2. Bell, A. M., Hankison, S. J. & Laskowski, K. L. The repeatability of behaviour: a meta-analysis. Anim Behav 77, 771–783 (2009).

3. Webster, M. M. & Ward, A. J. W. Personality and social context. Biol Rev 86, 759–773 (2011).

4. Wolf, M. & Krause, J. Why personality differences matter for social functioning and social structure. Trends Ecol Evol 29, 306–308 (2014).

5. Ward, A. J. W., Thomas, P., Hart, P. J. B. & Krause, J. Correlates of boldness in three-spined sticklebacks (*Gasterosteus aculeatus*). Behav Ecol Sociobiol 55, 561–568 (2004).

6. Kurvers, R. H. J. M. et al. Personality differences explain leadership in barnacle geese. Anim Behav 78, 447–453 (2009).

7. Harcourt, J. L., Ang, T. Z., Sweetman, G., Johnstone, R. A. & Manica, A. Social feedback and the emergence of leaders and followers. Curr Bio 19, 248–252 (2009).

8. Pettit, B., Ákos, Z., Vicsek, T. & Biro, D. Speed determines leadership and leadership determines learning during pigeon flocking. Curr Bio 25, 3132–3137 (2015).

9. Pike, T. W., Samanta, M., Lindström, J. & Royle, N. J. Behavioural phenotype affects social interactions in an animal network. Proc Biol Sci 275, 2515–2520 (2008).

10. Aplin, L. M. et al. Individual personalities predict social behaviour in wild networks of great tits (*Parus major*). Ecol Lett 16, 1365–72 (2013).

11. Jolles, J. W. et al. The role of social attraction and its link with boldness in the collective movements of three-spined sticklebacks. Animal Behaviour 99, 147–153 (2015).

12. Farine, D., Strandburg-Peshkin, A., Couzin, I., Berger-Wolf, T. & Crofoot, M. Individual variation in local interaction rules can explain emergent patterns of spatial organisation in wild baboons. Proc Biol Sci 25–29 (2017).

13. Dyer, J. R. G., Croft, D. P., Morrell, L. J. & Krause, J. Shoal composition determines foraging success in the guppy. Behav Ecol 20, 165–171 (2009).

14. Laskowski, K. L., Montiglio, P.-O. & Pruitt, J. N. Individual and group performance suffers from social niche disruption. Am Nat 187, 766–785 (2016).

15. Pruitt, J. N. & Riechert, S. E. How within-group behavioural variation and task efficiency enhance fitness in a social group. Proc Biol Sci 278, 1209–1215 (2011).

16. Ioannou, C. C. & Dall, S. R. X. Individuals that are consistent in risk-taking benefit during collective foraging. Sci Rep 6, 33991 (2016).

17. Smith, B. R. & Blumstein, D. T. Fitness consequences of personality: a meta-analysis. Behav Ecol 19, 448–455 (2008).

18. Sih, A., Cote, J., Evans, M., Fogarty, S. & Pruitt, J. N. Ecological implications of behavioural syndromes. Ecol Lett 15, 278–289 (2012).

19. Réale, D., Dingemanse, N. J., Kazem, A. J. N. & Wright, J. Evolutionary and ecological approaches to the study of personality. Phil Trans R Soc B 365, 3937–3946 (2010).

20. Schaerf, T. M., Herbert-Read, J. E., Myerscough, M. R., Sumpter, D. J. T. & Ward, A. J. W. Identifying differences in the rules of interaction between individuals in moving animal groups. arXiv preprint arXiv:1601.08202 (2016). 1601.08202.

21. Couzin, I. D. & Krause, J. Self-organization and collective behavior in vertebrates. Adv Stud Behav 32, 1–75 (2003).

22. Sumpter, D. J. T. Collective animal behavior (Princeton University Press, Oxford, 2010).

23. Herbert-Read, J. E. Understanding how animal groups achieve coordinated movement. J Exp Biol 219, 2971–2983 (2016).

24. Couzin, I. D., Krause, J., James, R., Ruxton, G. D. & Franks, N. R. Collective memory and spatial sorting in animal groups. J Theor Biol 218, 1–11 (2002).

25. Tunstrøm, K. et al. Collective states, multistability and transitional behavior in schooling fish. PLoS Comput Biol 9, e1002915 (2013).

26. Couzin, I. D., Krause, J., Franks, N. R. & Levin, S. A. Effective leadership and decision-making in animal groups on the move. Nature 433, 513–516 (2005).

27. Ioannou, C. C., Singh, M. & Couzin, I. D. Potential leaders trade off goal-oriented and socially oriented behavior in mobile animal groups. Am Nat 186, 284–293 (2015).

28. Persson, L. & Eklöv, P. Prey refuges affecting interactions between piscivorous Perch and juvenile Perch and Roach. Ecology 76, 70–81 (1995).

29. Jolles, J. W., Manica, A. & Boogert, N. J. Food intake rates of inactive fish are positively linked to boldness in three-spined sticklebacks *Gasterosteus aculeatus*. J Fish Biol 88, 1661–1668 (2016).

30. Krause, J. & Ruxton, G. D. Living in groups (Oxford University Press, Oxford, 2002).

31. Ward, A. & Webster, M. Sociality: the behaviour of group-living animals (Springer International Publishing, Switzerland, 2016).

32. Katz, Y., Tunstrøm, K., Ioannou, C. C., Huepe, C. & Couzin, I. D. Inferring the structure and dynamics of interactions in schooling fish. Proc Natl Acad Sci 108, 18720–18725 (2011).

33. Herbert-Read, J. E. et al. Inferring the rules of interaction of shoaling fish. Proc Natl Acad Sci 108, 18726–18731 (2011).

34. Herbert-Read, J. E. et al. The role of individuality in collective group movement. Proc Biol Sci 280, 20122564 (2012).

35. Nagy, M., Akos, Z., Biro, D. & Vicsek, T. Hierarchical group dynamics in pigeon flocks. Nature 464, 890–893 (2010).

36. Johnstone, R. A. & Manica, A. Evolution of personality differences in leadership. Proc Natl Acad Sci 108, 8373–8378 (2011).

37. Farine, D. R., Montiglio, P.-o. & Spiegel, O. From individuals to groups and back: the evolutionary implications of group phenotypic composition. Trends Ecol Evol 30, 609–621 (2015).

38. Spiegel, O., Leu, S. T., Bull, C. M. & Sih, A. What’s your move? Movement as a link between personality and spatial dynamics in animal populations. Ecol Lett 20, 3–18 (2017).

39. Borg, B., Bornestaf, C., Hellqvist, A., Schmitz, M. & Mayer, I. Mechanisms in the photoperiodic control of reproduction in the Stickleback. Behav 141, 1521–1530 (2004).

40. Jolles, J. W., Aaron Taylor, B. & Manica, A. Recent social conditions affect boldness repeatability in individual sticklebacks. Anim Behav 112, 139–145 (2016).

41. Biro, P. A. Do rapid assays predict repeatability in labile (behavioural) traits? Anim Behav 83, 1295–1300 (2012).

42. Webster, M. M. & Laland, K. N. Evaluation of a non-invasive tagging system for laboratory studies using three-spined sticklebacks *Gasterosteus aculeatus*. J Fish Biol 75, 1868–1873 (2009).

43. Wright, D. & Krause, J. Repeated measures of shoaling tendency in zebrafish (*Danio rerio*) and other small teleost fishes. Nat Protoc 1, 1828–1831 (2006).

44. Crawley, M. The R book (Wiley, Chichester, 2007).

45. Nakagawa, S. & Schielzeth, H. Repeatability for Gaussian and non-Gaussian data: a practical guide for biologists. Biol Rev 85, 935–956 (2010).

